# Impact of vancomycin loading doses and dose escalation on glomerular function and kidney injury biomarkers in a translational rat model

**DOI:** 10.1101/2022.09.26.509628

**Authors:** Jack Chang, Gwendolyn M. Pais, Patti L. Engel, Patryk Klimek, Sylwia Marianski, Kimberly Valdez, Marc H. Scheetz

## Abstract

Vancomycin induced kidney injury is common, and outcomes in humans are well predicted by animal models. This study employed our translational rat model to investigate temporal changes in glomerular filtration rate (GFR) and correlation with kidney injury biomarkers related to various vancomycin dosing strategies. First, Sprague Dawley rats received allometrically scaled loading doses or standard doses. Rats that received a loading dose had lower GFR and increased urinary injury biomarkers (kidney injury molecule 1 [KIM-1] and clusterin) that persisted through day 2, compared to those that did not receive a loading dose. Second, we compared low and high allometrically scaled vancomycin doses to a positive acute kidney injury control of high dose folic acid. Rats in both the low and high vancomycin dose groups had higher GFRs on all dosing days versus the positive control group. When the two vancomycin groups were compared, rats that received the low dose had significantly higher GFR on days 1, 2, and 4. Compared to low dose vancomycin, KIM-1 was elevated in high dose rats on dosing day 3. GFR correlated most closely with the urinary injury biomarker KIM-1, on all experimental days.

Vancomycin loading doses were associated with significant loss of kidney function and elevation of urinary injury biomarkers. In our translational rat model, both the degree of kidney function decline and urinary biomarker rise corresponded to the magnitude of vancomycin dose (i.e. higher dose resulted in more kidney function decline and greater degree of urinary injury biomarker increase).

## Introduction

Vancomycin is a glycopeptide antibiotic that remains the treatment of choice for methicillin-resistant *Staphylococcus aureus* (MRSA) infections. Unfortunately, vancomycin-induced kidney injury is a common adverse effect which occurs at an attributable rate of at least 10% (1). In the current guidelines for vancomycin dosing in *Staphylococcus aureus* infections, loading doses are recommended for patients who are critically ill, require dialysis or renal replacement therapy, have serious MRSA infections, or are receiving vancomycin continuous infusion therapy (2). However, it is unknown whether vancomycin loading doses contribute to increased kidney injury. Clinical studies that underlie this recommendation assume increased efficacy and are mostly retrospective, contain few patients, and are based on previous trough goals for vancomycin monitoring (3-6). As a result, there exists a need to investigate the nephrotoxic potential of vancomycin loading doses using sensitive and specific biomarkers for kidney function and injury. The goals of this study were thus two-fold. First, we examined kidney injury and function in the setting of a loading dose and that of no loading dose. Second, we defined the dose:response relationship of standard dose and high dose vancomycin to a positive control of acute kidney injury. To do so, we employed our translational rat model, using iohexol clearance as a surrogate for kidney function, and urinary kidney injury biomarkers (kidney injury molecule-1 [KIM-1], osteopontin, and clusterin) (7-10). Our previous work using this model has shown that exposures and outcomes relating to vancomycin-induced kidney injury are well predicted by the rat, and closely link to prospective human studies (11). In addition, increases in specific urinary biomarkers of kidney injury precede declines in glomerular filtration rate (GFR) in response to nephrotoxins (10, 12). The goal of this study was to investigate temporal changes in glomerular function and urinary injury biomarkers related to variable vancomycin doses.

## Results

### Characteristics of animal cohort

A total of 34 male Sprague Dawley rats were assigned to two separate experimental arms (1: investigation of the impact of vancomycin loading doses, 2: investigation of variable vancomycin doses with an AKI-positive control), with dosing group assignments shown in Supplementary table 1. One animal provided only terminal plasma samples due to occluded catheters; all other animals contributed complete data. The mean baseline weight of the rats was 274.9 g with a standard deviation (SD) of 9.4 g. Mean weight change was not significantly different over all experimental days, among rats in the loading dose experiment. In the AKI-positive control experiment, rats in the folic acid (AKI-positive control) group experienced significant mean weight loss by day 3, when compared to rats in the low and high vancomycin dose groups (-26.8 g vs. -1.2 g vs. -4.0 g, p<0.0001).

**Table 1:** Marginal differences versus VAN no load group as referent.

| **Table 1: Marginal differences versus VAN no load group as referent** |  |
| --- | --- |
| **Dosing day** | **Treatment groups** |
|  | **VAN loading dose group** |
| GFR difference (mL/min/100 g B.W.), mean (95% CI), p-value |  |
| Baseline | -0.01 (-0.09 to 0.07), p=0.79 |
| Day 1-AM | **-0.16 (-0.24 to -0.08), p<0.001** |
| Day 1-PM | **-0.12 (-0.19 to -0.04), p=0.005** |
| Day 2 | **-0.11 (-0.19 to -0.03), p=0.007** |
| Day 3 | -0.07 (-0.15 to 0.01), p=0.08 |
| Day 4 | -0.06 (-0.14 to 0.03), p=0.18 |
| Urinary KIM-1 difference (ng), mean (95% CI), p-value |  |
| Baseline | 0.78 (-46.9 to 48.5), p=0.97 |
| Day 1-AM | **106.9 (59.3 to 154.6), p<0.001** |
| Day 1-PM | 35.9 (-11.8 to 83.6), p=0.14 |
| Day 2 | **81.8 (34.1 to 129.5), p=0.001** |
| Day 3 | 25.4 (-22.3 to 73.0), p=0.29 |
| Day 4 | 15.0 (-32.7 to 62.7), p=0.54 |
| Urinary clusterin difference (ng), mean (95% CI), p-value |  |
| Baseline | 3536 (-1486 to 8559), p=0.17 |
| Day 1-AM | **8247 (3224 to 13269), p=0.001** |
| Day 1-PM | 2553 (-2469 to 7576), p=0.32 |
| Day 2 | **7612 (2485 to 12740), p=0.004** |
| Day 3 | 4159 (-864 to 9181), p=0.11 |
| Day 4 | 2917 (-2105 to 7940), p=0.26 |
| Urinary OPN difference (ng), mean (95% CI), p-value |  |
| Baseline | -0.15 (-3.96 to 3.66), p=0.94 |
| Day 1-AM | **6.24 (3.06 to 9.42), p<0.001** |
| Day 1-PM | 2.36 (-0.82 to 5.54), p=0.15 |
| Day 2 | 2.40 (-0.86 to 5.66), p=0.15 |
| Day 3 | 0.29 (-2.89 to 3.46), p=0.86 |
| Day 4 | -0.84 (-4.02 to 2.34), p=0.61 |
| Urine output difference (mL), mean (95% CI), p-value |  |
| Baseline | 3.86 (-0.07 to 7.79), p=0.054 |
| Day 1-AM | 3.31 (-0.63 to 7.24), p=0.09 |
| Day 1-PM | 2.13 (-1.80 to 6.06), p=0.29 |
| Day 2 | 3.38 (-0.55 to 7.31), p=0.09 |
| Day 3 | **5.11 (1.18 to 9.04), p=0.01** |
| Day 4 | 0.43 (-3.51 to 4.36), p=0.83 |

### Kidney function change over time

Baseline (i.e. prior to therapy) mean GFR was not significantly different between rats enrolled into the vancomycin (VAN) no load and VAN loading dose groups (0.45 [95% CI: 0.42 to 0.49] vs. 0.46 [95% CI: 0.41 to 0.50] mL/min/100 g body weight, p=0.76). After receipt of the VAN loading dose in the morning of day 1, rats in the VAN loading dose group experienced a significant decline in GFR compared to the VAN no loading dose group (Figure 1: -0.16 mL/min/100 g body weight, 95% CI: -0.24 to -0.08, p<0.001). A significantly lower GFR was also observed among the VAN loading dose group rats after receipt of the maintenance VAN dose in the evening of day 1 (-0.12 mL/min/100 g body weight, 95% CI: -0.19 to -0.04, p=0.005), which persisted through day 2 (-0.11 mL/min/100 g body weight, 95% CI: -0.19 to -0.03, p=0.007).

**Figure 1:**
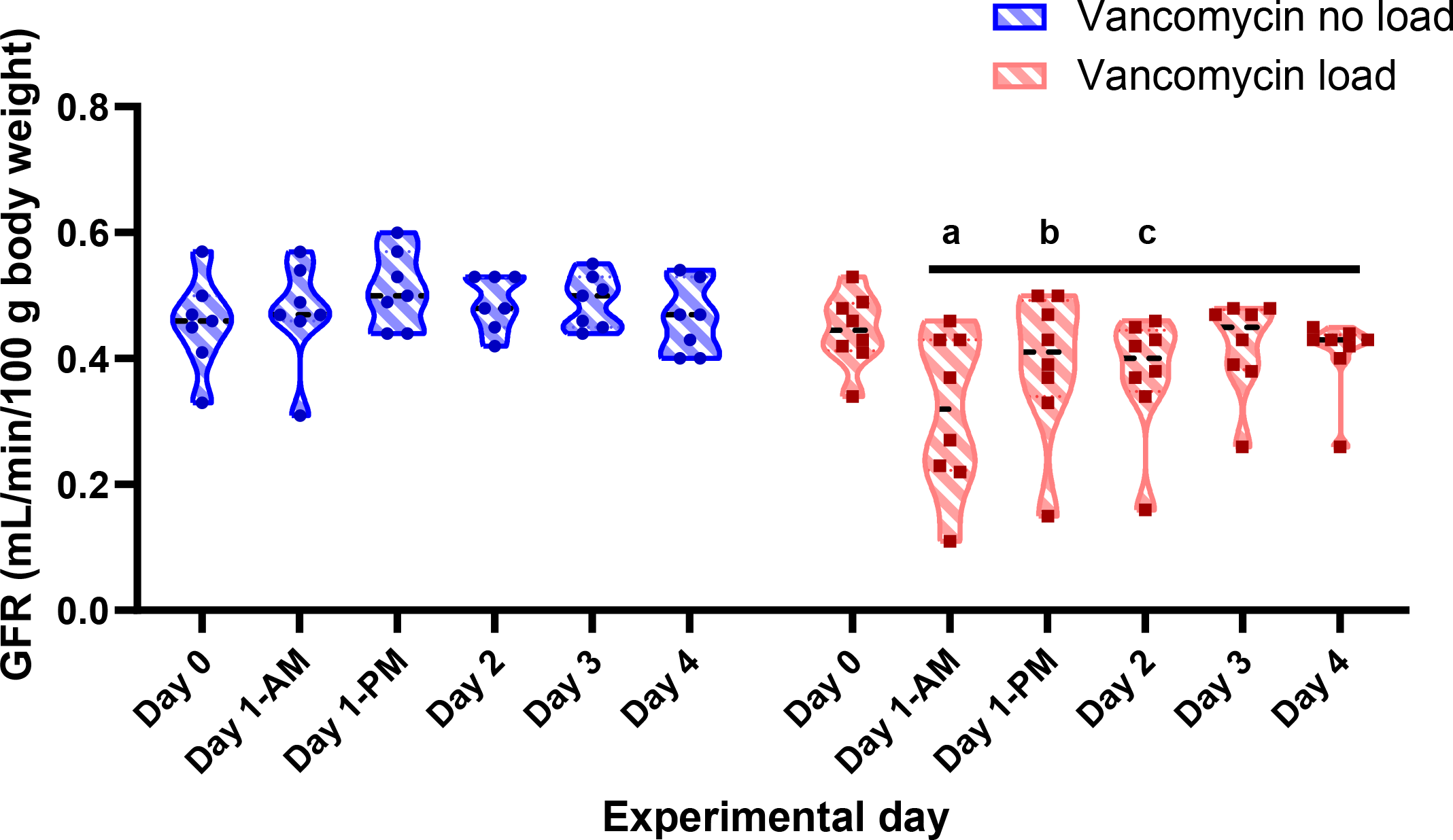
Iohexol GFR comparison for rats in the loading dose arm, between treatment groups and across dosing days. Comparing the treatment groups by experimental day, GFR was significantly lower on day 1 in the morning (a: -0.16 mL/min/100g body weight, 95% CI: -0.24 to -0.08, p<0.001), day 1 in the evening (b: -0.12 mL/min/100g body weight, 95% CI: -0.19 to -0.04, p=0.005), and day 2 (c: -0.11 mL/min/100g body weight, 95% CI: -0.19 to - 0.03, p=0.007) for rats that received a vancomycin loading dose. No significant changes in GFR from baseline (day 0) were identified among rats that did not receive a vancomycin loading dose.

In the second experimental arm, baseline GFR (i.e. prior to drug administration) was not significantly different for rats that were assigned to any of the treatment groups; folic acid (AKI-positive control), low VAN dose, and high VAN dose groups (0.42 [95% CI: 0.45 to 0.50], 0.43 [95% CI: 0.43 to 0.51], 0.43 [95% CI: 0.39 to 0.52] mL/min/100 g body weight, respectively p=0.99). Compared to the AKI-positive control group, GFR was significantly higher among rats in the low VAN dose group on day 1 (Figure 2: 0.33 mL/min/100 g body weight, 95% CI: 0.22 to 0.44, p<0.001), day 2 (0.34 mL/min/100 g body weight, 95% CI: 0.23 to 0.45, p<0.001), day 3 (0.29 mL/min/100 g body weight, 95% CI: 0.18 to 0.40, p<0.001), and day 4 (0.39 mL/min/100 g body weight, 95% CI: 0.29 to 0.51, p<0.001). Compared to the AKI-positive control group, GFR was also significantly higher among rats in the high VAN dose group on day 1 (0.21 mL/min/100 g body weight, 95% CI: 0.09 to 0.32, p<0.001), day 2 (0.22 mL/min/100 g body weight, 95% CI: 0.11 to 0.33, p<0.001), day 3 (0.24 mL/min/100 g body weight, 95% CI: 0.13 to 0.35, p<0.001), and day 4 (0.26 mL/min/100 g body weight, 95% CI: 0.15 to 0.37, p<0.001). In a direct comparison of the low and high VAN dose groups, GFR was significantly higher among rats in the low VAN dose group on day 1 (0.13 mL/min/100 g body weight, 95% CI: 0.02 to 0.24, p=0.025), day 2 (0.12 mL/min/100 g body weight, 95% CI: 0.01 to 0.23, p=0.037), and day 4 (0.14 mL/min/100 g body weight, 95% CI: 0.03 to 0.25, p=0.012).

**Figure 2:**
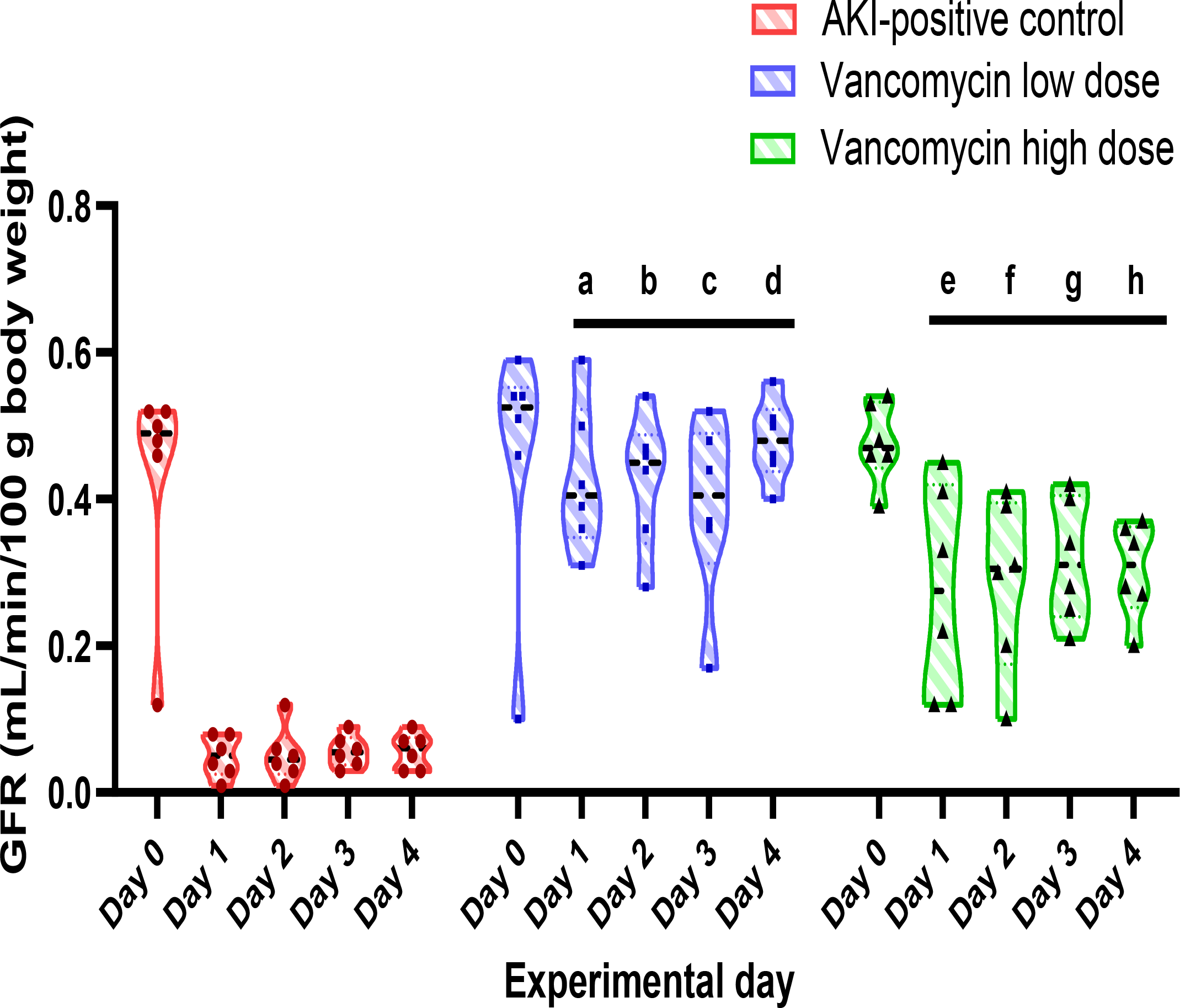
Iohexol GFR comparison for rats in the variable vancomycin dose arm, between treatment groups and across dosing days. Using the AKI-positive control as the referent group, GFR was significantly higher in both the low and high VAN dose group rats, on all dosing days (a: 0.33 mL/min/100g body weight, 95% CI: 0.22 to 0.44, p<0.001, b: 0.34 mL/min/100 g body weight, 95% CI: 0.23 to 0.45, p<0.001, c: 0.29 mL/min/100 g body weight, 95% CI: 0.18 to 0.40, p<0.001, d: 0.39 mL/min/100 g body weight, 95% CI: 0.29 to 0.51, p<0.001, e: 0.21 mL/min/100 g body weight, 95% CI: 0.09 to 0.32, p<0.001, f: 0.22 mL/min/100 g body weight, 95% CI: 0.11 to 0.33, p<0.001, g: 0.24 mL/min/100 g body weight, 95% CI: 0.13 to 0.35, p<0.001, h: 0.26 mL/min/100 g body weight, 95% CI: 0.15 to 0.37, p<0.001).

### Urine output and injury biomarkers

Baseline urine output (prior to drug administration) was not significantly different between the treatment groups of the loading dose arm (Supplementary table 1: 8.9 mL/24 hours, 95% CI: 5.7 to 12.4 vs. 12.7 mL/24 hours, 95% CI: 6.5 to 18.0, p=0.054). Urine output was significantly higher on day 3 among rats that received a VAN loading dose (Table 1: 5.1 mL/24 hours, 95% CI: 1.2 to 9.0, p=0.01).

Compared to baseline, rats in the VAN loading dose group showed significantly elevated levels of urinary kidney injury molecule-1 (KIM-1) after receipt of the VAN loading dose in the morning of day 1 [day 1-AM] (Figure 3a: 106.9 ng/24 hours, 95% CI: 59.3 to 154.6, p<0.001), and day 2 (81.8 ng/24 hours, 95% CI: 34.1 to 129.5, p=0.001). Rats that received a VAN loading dose also had significantly elevated levels of urinary clusterin on day 1-AM (Figure 3b: 8247 ng/24 hours, 95% CI: 3224 to 13269, p=0.001) and day 2 (7612 ng/24 hours, 95% CI: 2485 to 12740, p=0.004). Significant elevations in urinary osteopontin (OPN) were observed after receipt of the VAN loading dose on day 1-AM (Figure 3c: 6.24 ng/24 hours, 95% CI: 3.06 to 9.42, p<0.001).

**Figure 3:**
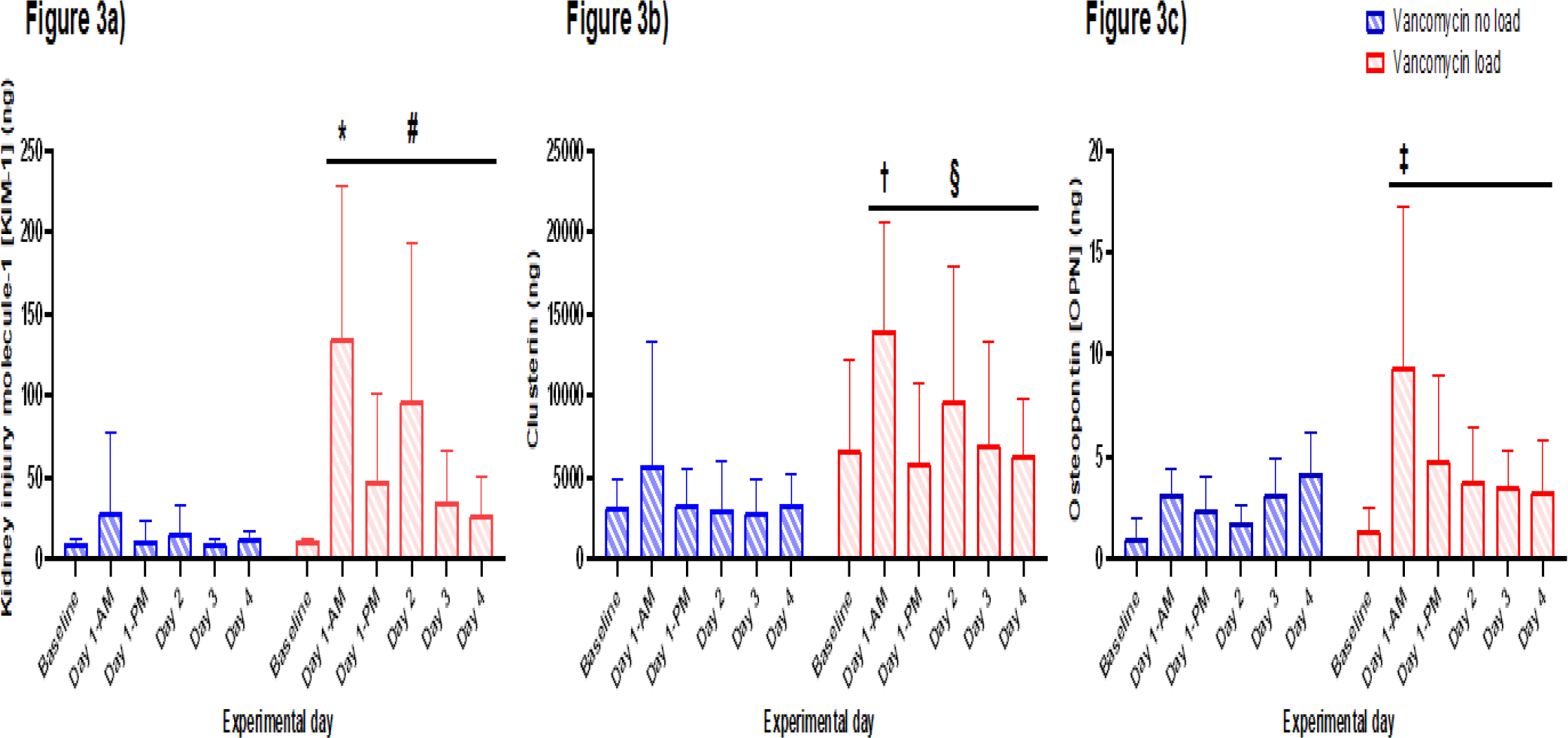
Comparison of urinary injury biomarker levels for rats in the loading dose arm, between treatment groups and across dosing days. The vancomycin no load group was used as the comparator. Significant differences in urinary KIM-1 were observed on day 1-AM (Figure 3a; *: 106.9 ng/24 hours, 95% CI: 59.3 to 154.6, p<0.001), and day 2 (#: 81.8 ng/24 hours, 95% CI: 34.1 to 129.5, p=0.001) among rats in the vancomycin load group. Significant differences in urinary clusterin were observed on day 1-AM (Figure 3b; †: 8247 ng/24 hours, 95% CI: 3224 to 13269, p=0.001) and day 2 (§: 7612 ng/24 hours, 95% CI: 2485 to 12740, p=0.004). A significant difference in urinary OPN was observed on day 1-AM (Figure 3c; ‡: 6.24 ng/24 hours, 95% CI: 3.06 to 9.42, p<0.001).

In the investigation of variable vancomycin doses with an AKI-positive control arm, baseline urine output before treatment was not significantly different between the treatment groups (Supplementary table 2: 8.9 mL/24 hours, 95% CI: 7.0 to 10.0 vs. 12.6 mL/24 hours, 95% CI: 9.3 to 12.6 vs. 11.3 mL/24 hours, 95% CI: 8.6 to 13.3, p=0.35). When compared to the group that received folic acid (AKI-positive control), rats in the low VAN dose group had significantly lower urine output on day 2 (Table 2: -16.1 mL/24 hours, 95% CI: -21.7 to -10.6, p<0.001) and day 3 (-10.3 mL/24 hours, 95% CI: -15.8 to -4.8, p<0.001). Rats in the high VAN dose group also had significantly lower urine output on day 2 (-10.3 mL/24 hours, 95% CI: -15.9 to -4.8, p<0.001) and day 3 (-9.8 mL/24 hours, 95% CI: -15.3 to -4.2, p=0.001), when compared to the group that received folic acid.

**Table 2:** Marginal differences versus AKI-positive control (folic acid) as referent group.

| Dosing day | Treatment groups |  |
| --- | --- | --- |
|  | Low VAN dose | High VAN dose |
| GFR difference (mL/min/100 g B.W.), mean (95% CI), p-value |  |  |
| Baseline (day 0) | 0.01 (-0.11 to 0.12), p=0.86 | 0.01 (-0.10 to 0.12), p=0.86 |
| Day 1 | **0.33 (0.22 to 0.44), p<0.001** | **0.21 (0.09 to 0.32), p<0.001** |
| Day 2 | **0.34 (0.23 to 0.45), p<0.001** | **0.22 (0.11 to 0.33), p<0.001** |
| Day 3 | **0.29 (0.18 to 0.40), p<0.001** | **0.24 (0.13 to 0.35), p<0.001** |
| Day 4 | **0.39 (0.29 to 0.51), p<0.001** | **0.26 (0.15 to 0.37), p<0.001** |
| Urinary KIM-1 difference (ng), mean (95% CI), p-value |  |  |
| Baseline | 0.27 (-74.7 to 75.3), p=0.99 | -0.88 (-75.9 to 74.1), p=0.98 |
| Day 1 | **-107.8 (-182.8 to -32.8), p=0.005** | -42.5 (-117.5 to 32.5), p=0.27 |
| Day 2 | **-172.0 (-247.0 to -97.1), p<0.001** | -74.6 (-149.6 to 0.46), p=0.051 |
| Day 3 | **-119.7 (-194.7 to -44.7), p=0.002** | **-77.2 (-152.2 to -2.16), p=0.044** |
| Day 4 | **-77.1 (-152.1 to -2.11), p=0.044** | 21.6 (-53.4 to 96.6), p=0.57 |
| Urinary clusterin difference (ng), mean (95% CI), p-value |  |  |
| Baseline | 113 (-7053 to 7280), p=0.98 | 170 (-6997 to 7336), p=0.96 |
| Day 1 | -6492 (-13658 to 675), p=0.08 | 23 (-7143 to 7189), p=0.99 |
| Day 2 | **-10326 (-17493 to -3159), p=0.005** | 2611 (-4556 to 9777), p=0.48 |
| Day 3 | -7010 (-14177 to 156), p=0.06 | 1029 (-6138 to 8195), p=0.78 |
| Day 4 | **-7841 (-15008 to -674), p=0.032** | **-8019 (-15509 to -528), p=0.036** |
| Urinary OPN difference (ng), mean (95% CI), p-value |  |  |
| Baseline | 0.16 (-9.81 to 10.1), p=0.98 | -0.35 (-11.5 to 10.8), p=0.95 |
| Day 1 | **-19.2 (-27.4 to -11.1), p<0.001** | **-18.5 (-26.3 to -10.7), p<0.001** |
| Day 2 | **-26.0 (-34.2 to -17.9), p<0.001** | **-24.3 (-32.1 to -16.5), p<0.001** |
| Day 3 | **-16.9 (-25.6 to -8.35), p<0.001** | **-18.5 (-26.3 to -10.7), p<0.001** |
| Day 4 | **-28.8 (-36.6 to -21.0), p<0.001** | **-30.1 (-39.4 to -20.8), p<0.001** |
| Urine output difference (mL), mean (95% CI), p-value |  |  |
| Baseline | 3.7 (-1.8 to 9.2), p=0.19 | 2.4 (-3.2 to 7.9), p=0.39 |
| Day 1 | -2.9 (-8.4 to 2.7), p=0.31 | -2.4 (-7.9 to 3.1), p=0.39 |
| Day 2 | **-16.1 (-21.7 to -10.6), p<0.001** | **-10.3 (-15.9 to -4.8), p<0.001** |
| Day 3 | **-10.3 (-15.8 to -4.8), p<0.001** | **-9.8 (-15.3 to -4.2), p=0.001** |
| Day 4 | -0.9 (-6.5 to 4.6), p=0.75 | -0.1 (-5.7 to 5.4), p=0.97 |

Compared to the group which received folic acid (AKI-positive control), rats in the low VAN dose group had significantly lower urinary KIM-1 on day 1 (Figure 4a: -107.8 ng/24 hours, 95% CI: -182.8 to -32.8, p=0.005), day 2 (-172.0 ng/24 hours, 95% CI: -247.0 to -97.1, p<0.001), day 3 (-119.7 ng/24 hours, 95% CI: -194.7 to - 44.7, p=0.002), and day 4 (-77.1 ng/24 hours, 95% CI: -152.1 to -2.11, p=0.044). Rats in the low VAN dose group also had significantly lower urinary clusterin on day 2 (Figure 4b: -10326 ng/24 hours, 95% CI: -17493 to -3159, p=0.005) and day 4 (-7841 ng/24 hours, 95% CI: -15008 to -674, p=0.032). Urinary OPN was also significantly lower among the low VAN dose group rats on day 1 (-19.2 ng/24 hours, 95% CI: -27.4 to -11.1, p<0.001), day 2 (-26.0 ng/24 hours, 95% CI: -34.2 to -17.9, p<0.001), day 3 (-16.9 ng/24 hours, 95% CI: -25.6 to -8.35, p<0.001), and day 4 (-28.8 ng/24 hours, 95% CI: -36.6 to -21.0, p<0.001).

**Figure 4:**
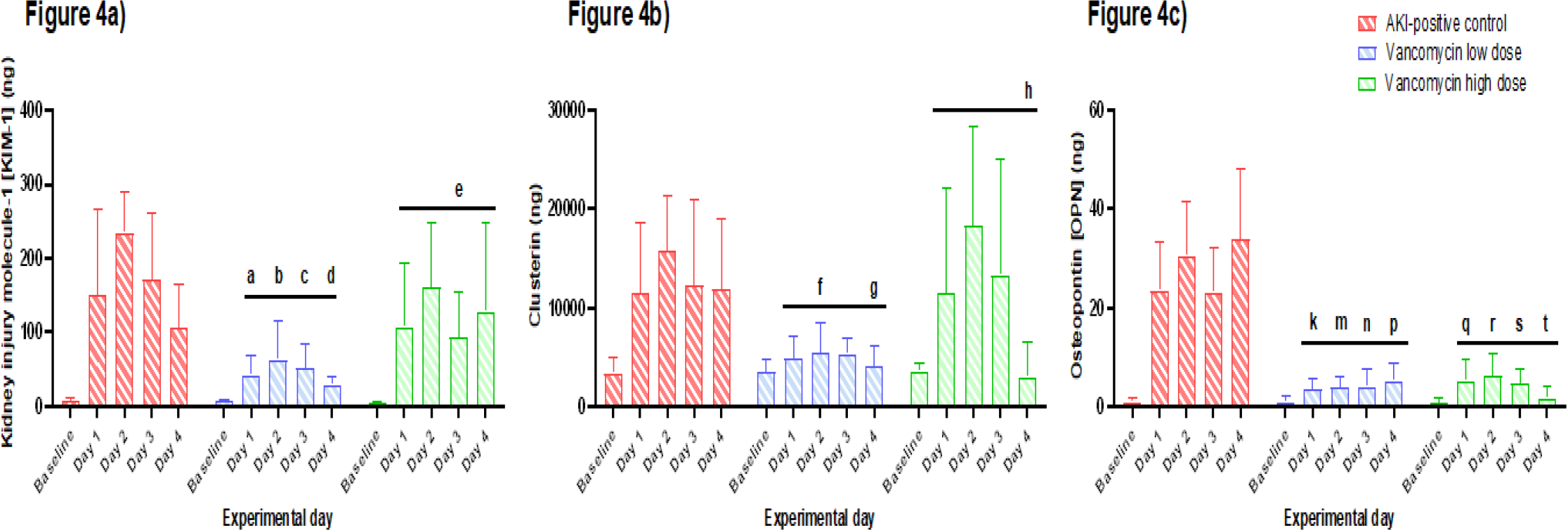
Comparison of urinary injury biomarker levels for rats in the variable vancomycin dose arm, between treatment groups and across dosing days. The folic acid group (AKI-positive control) was used as the comparator. Among rats in the low VAN group, significant differences in urinary KIM-1 were observed on day 1 (Figure 4a; a: -107.8 ng/24 hours, 95% CI: -182.8 to -32.8, p=0.005), day 2 (b: -172.0 ng/24 hours, 95% CI: -247.0 to -97.1, p<0.001), day 3 (c: -119.7 ng/24 hours, 95% CI: -194.7 to -44.7, p=0.002), and day 4 (d: -77.1 ng/24 hours, 95% CI: -152.1 to -2.11, p=0.044). Urinary clusterin was significantly lower on day 2 (Figure 4b; f: - 10326 ng/24 hours, 95% CI: -17493 to -3159, p=0.005) and day 4 (g: -7841 ng/24 hours, 95% CI: -15008 to - 674, p=0.032). Urinary OPN was significantly lower on day 1 (Figure 4c; k: -19.2 ng/24 hours, 95% CI: -27.4 to - 11.1, p<0.001), day 2 (m: -26.0 ng/24 hours, 95% CI: -34.2 to -17.9, p<0.001), day 3 (n: -16.9 ng/24 hours, 95% CI: -25.6 to -8.35, p<0.001), and day 4 (p: -28.8 ng/24 hours, 95% CI: -36.6 to -21.0, p<0.001). Among rats in the high VAN group, significant differences in urinary KIM-1 were observed on day 3 (e: -77.2 ng/24 hours, 95% CI: -152.2 to -2.16, p=0.044). Urinary clusterin was significantly lower on day 4 (Figure 4b; h: -8019 ng/24 hours, 95% CI: -15509 to -528, p=0.036). Urinary OPN was significantly lower on day 1 (Figure 4c; q: -18.5 ng/24 hours, 95% CI: -26.3 to -10.7, p<0.001), day 2 (r: -24.3 ng/24 hours, 95% CI: -32.1 to -16.5, p<0.001), day 3 (s: -18.5 ng/24 hours, 95% CI: -26.3 to -10.7, p<0.001), and day 4 (t: -30.1 ng/24 hours, 95% CI: -39.4 to -20.8, p<0.001).

Compared to the folic acid group, rats in the high VAN dose group had significantly lower urinary KIM-1 on day 3 (Figure 4a: -77.2 ng/24 hours, 95% CI: -152.2 to -2.16, p=0.044). Rats in the high VAN dose group had significantly lower urinary clusterin on day 4 (-8019 ng/24 hours, 95% CI: -15509 to -528, p=0.036). Urinary OPN was also significantly lower among the high VAN dose group rats on day 1 (-18.5 ng/24 hours, 95% CI: - 26.3 to -10.7, p<0.001), day 2 (-24.3 ng/24 hours, 95% CI: -32.1 to -16.5, p<0.001), day 3 (-18.5 ng/24 hours, 95% CI: -26.3 to -10.7, p<0.001), and day 4 (-30.1 ng/24 hours, 95% CI: -39.4 to -20.8, p<0.001).

In a direct comparison of the low and high VAN dose groups, rats in the low VAN dose group had significantly lower urinary KIM-1 on day 2 (-97.5 ng/24 hours, 95% CI: -165.9 to -29.0, p=0.005) and day 4 (-98.8 ng/24 hours, 95% CI: -167.3 to -30.3, p=0.005). Rats in the low VAN dose group also had significantly lower urinary clusterin on day 2 (-12936 ng/24 hours, 95% CI: -20103 to -5770, p<0.001) and day 3 (-8039 ng/24 hours, 95% CI: -15206 to -873, p=0.028). No significant differences in urinary clusterin were seen between the low and high VAN dose groups.

### Correlation between urinary injury biomarkers and GFR

Spearman’s rank correlations between GFR and urinary kidney injury biomarkers in the loading dose experiment are listed in Table 3. Among rats that received a vancomycin loading dose, urinary KIM-1 was significantly correlated with decreasing GFR on day 1-AM (Figure 5a; Spearman’s rho: -0.94, p<0.0001), day 1-PM (Spearman’s rho: -0.66, p=0.008), day 2 (Spearman’s rho: -0.72, p=0.002), day 3 (Spearman’s rho: -0.53, p=0.042), and day 4 (Spearman’s rho: -0.66, p=0.007). Urinary clusterin was significantly correlated with decreasing GFR on day-1 AM (Figure 5b; Spearman’s rho: -0.89, p<0.0001), day 2 (Spearman’s rho: -0.75, p=0.002), and day 4 (Spearman’s rho: -0.78, p=0.0006). Urinary OPN was significantly correlated with decreasing GFR on day 1-AM (Figure 5c; Spearman’s rho: -0.64, p=0.01) and day 2 (Spearman’s rho: -0.59, p=0.03).

**Table 3:** Summary of urinary biomarker correlations with GFR in loading dose experiment.

|  |  |  |  |  |  |  |  |
| --- | --- | --- | --- | --- | --- | --- | --- |
| **Urinary biomarker** |  | **Baseline (day 0)** | **Day 1-AM** | **Day 1-PM** | **Day 2** | **Day 3** | **Day 4** |
| KIM-1 | Spearman's Rho | 0.05 | -0.94 | -0.66 | -0.72 | -0.53 | -0.66 |
|  | p-value | 0.86 | **<0.0001** | **0.008** | **0.002** | **0.042** | **0.007** |
| Clusterin | Spearman's Rho | 0.01 | -0.89 | -0.42 | -0.75 | -0.28 | -0.78 |
|  | p-value | 0.97 | **<0.0001** | 0.12 | **0.002** | 0.30 | **0.0006** |
| OPN | Spearman's Rho | 0.53 | -0.64 | -0.34 | -0.59 | -0.06 | -0.22 |
|  | p-value | 0.09 | **0.01** | 0.22 | **0.03** | 0.82 | 0.43 |

**Figure 5:**
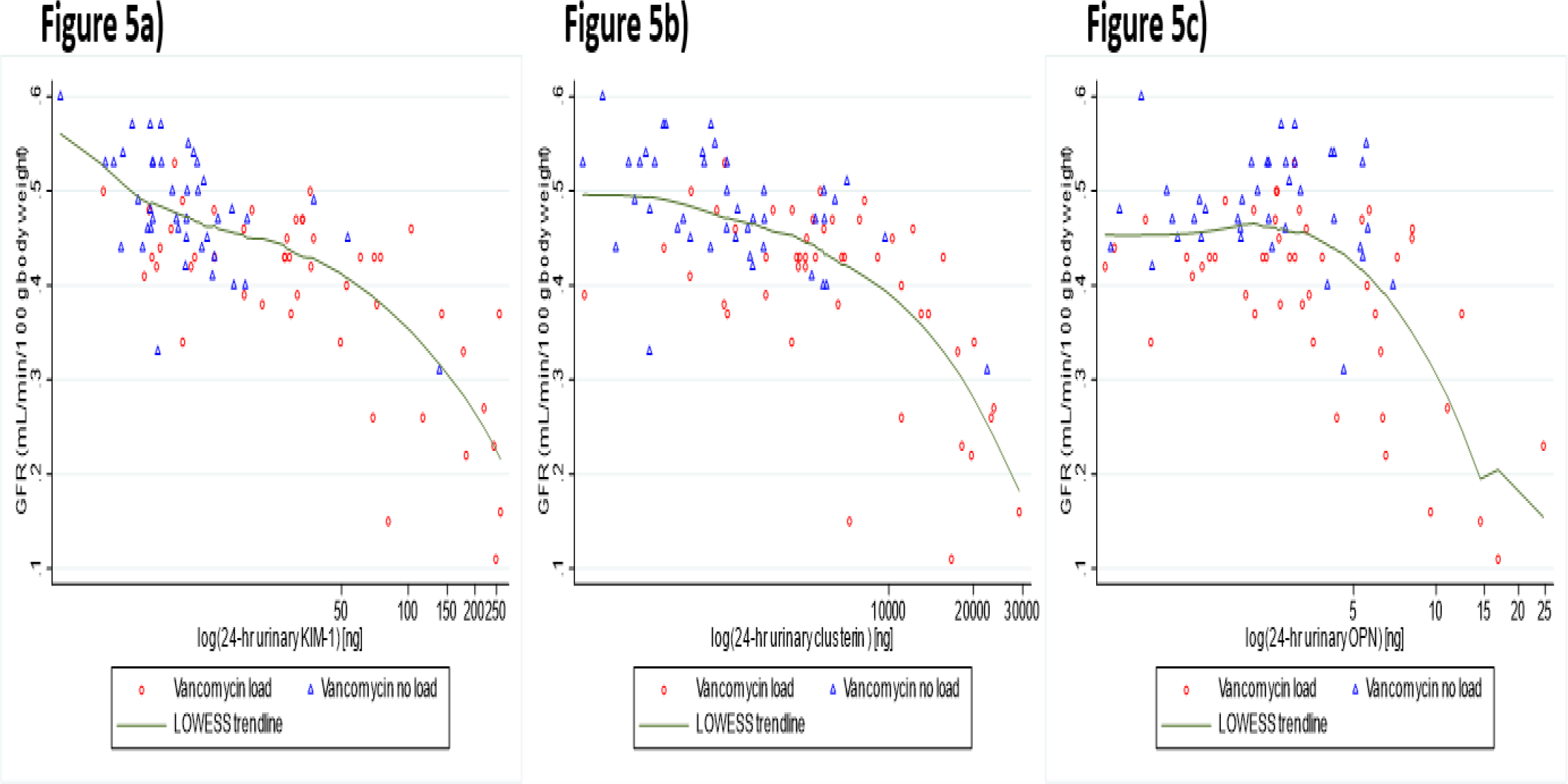
Spearman correlations of GFR with 24-hour urinary injury biomarker levels (biomarker amounts shown on logarithmic scale). Among rats which received a vancomycin loading dose (red data points), urinary KIM-1 was significantly correlated with decreasing GFR on day 1-AM (Figure 5a; Spearman’s rho: -0.94, p<0.0001), day 1-PM (Spearman’s rho: -0.66, p=0.008), day 2 (Spearman’s rho: -0.72, p=0.002), day 3 (Spearman’s rho: -0.53, p=0.042), and day 4 (Spearman’s rho: -0.66, p=0.007). Urinary clusterin was significantly correlated with decreasing GFR on day-1 AM (Figure 5b; Spearman’s rho: -0.89, p<0.0001), day 2 (Spearman’s rho: -0.75, p=0.002), and day 4 (Spearman’s rho: -0.78, p=0.0006). Urinary OPN was significantly correlated with decreasing GFR on day 1-AM (Figure 5c; Spearman’s rho: -0.64, p=0.01) and day 2 (Spearman’s rho: -0.59, p=0.03).

Spearman’s rank correlations between GFR and urinary kidney injury biomarkers in Aim 2 are listed in Table 4. Among rats in the AKI-positive control group, urinary KIM-1 was significantly correlated with decreasing GFR on day 1 (Figure 6a; Spearman’s rho: -0.56, p=0.016), day 2 (Spearman’s rho: -0.73, p=0.0006), day 3 (Spearman’s rho: -0.52, p=0.03), and day 4 (Spearman’s rho: -0.63, p=0.005). Urinary clusterin was significantly correlated with decreasing GFR on day 2 (Figure 6b; Spearman’s rho: -0.53, p=0.0248). Urinary OPN was significantly correlated with decreasing GFR on day 1 (Figure 6c; Spearman’s rho: -0.75, p=0.0005), day 2 (Spearman’s rho: -0.85, p<0.0001), day 3 (Spearman’s rho: -0.69, p=0.003), and day 4 (Spearman’s rho: -0.63, p=0.012).

**Table 4:** Summary of urinary biomarker correlations with GFR in AKI-positive control experiment.

|  |  |  |  |  |  |  |
| --- | --- | --- | --- | --- | --- | --- |
| Urinary biomarker |  | Baseline | Day 1 | Day 2 | Day 3 | Day 4 |
| KIM-1 | Spearman's Rho | -0.01 | -0.56 | -0.73 | -0.52 | -0.63 |
|  | p-value | 0.99 | **0.016** | **0.0006** | **0.03** | **0.005** |
| Clusterin | Spearman's Rho | 0.21 | -0.37 | -0.53 | -0.18 | -0.31 |
|  | p-value | 0.41 | 0.13 | **0.0248** | 0.48 | 0.22 |
| OPN | Spearman's Rho | -0.32 | -0.75 | -0.85 | -0.69 | -0.63 |
|  | p-value | 0.40 | **0.0005** | **<0.0001** | **0.003** | **0.012** |

**Figure 6:**
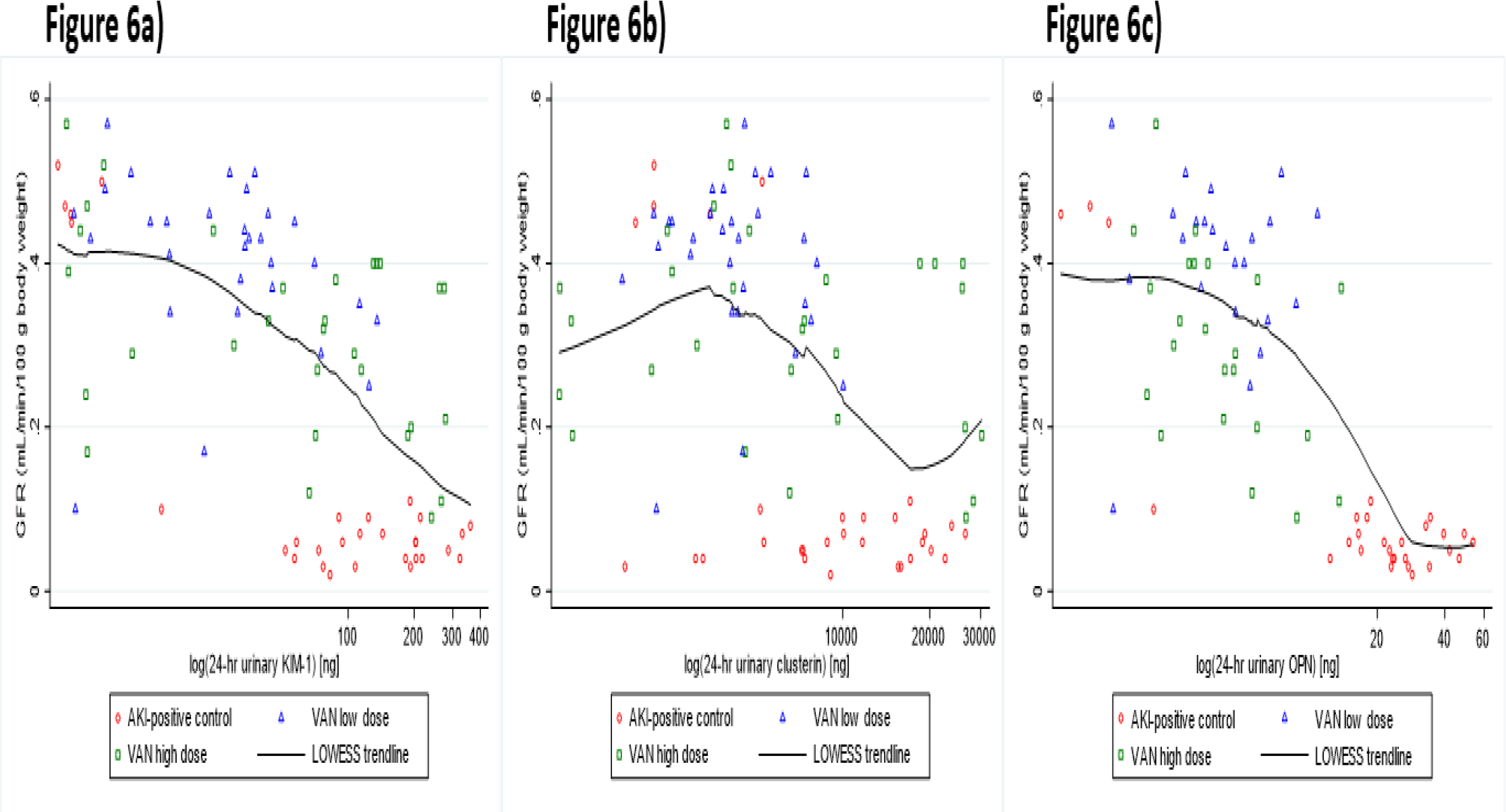
Spearman correlations of GFR with 24-hour urinary injury biomarker levels (biomarker amounts shown on logarithmic scale). Among rats in the AKI-positive control group (red data points), urinary KIM-1 was significantly correlated with decreasing GFR on day 1 (Figure 6a; Spearman’s rho: -0.56, p=0.016), day 2 (Spearman’s rho: -0.73, p=0.0006), day 3 (Spearman’s rho: -0.52, p=0.03), and day 4 (Spearman’s rho: -0.63, p=0.005). Urinary clusterin was significantly correlated with decreasing GFR on day 2 (Figure 6b; Spearman’s rho: -0.53, p=0.0248). Urinary OPN was significantly correlated with decreasing GFR on day 1 (Figure 6c; Spearman’s rho: -0.75, p=0.0005), day 2 (Spearman’s rho: -0.85, p<0.0001), day 3 (Spearman’s rho: -0.69, p=0.003), and day 4 (Spearman’s rho: -0.63, p=0.012).

## Discussion

In this study, we found that receipt of a vancomycin loading dose resulted in a significant decline in glomerular function (as assessed by iohexol clearance), immediately after dosing, which persisted for 24 hours until function was recovered to baseline levels in most rats. This represented a mean 31% relative drop from baseline before treatment, and from matched controls. Urinary injury biomarker changes paralleled functional changes; Significant rises in the urinary injury biomarkers KIM-1 and OPN occurred immediately after receipt of the vancomycin loading dose, which correlated with GFR declines immediately after receipt of the loading dose. Among the assessed urinary biomarkers, KIM-1 was most closely correlated with declining GFR over the course of the experiment. Notably, neither significant changes in GFR nor significant elevations in urinary injury biomarkers were observed among rats that did not receive a vancomycin loading dose. These findings of increased kidney injury and loss of function in response to receipt of a vancomycin loading dose are both novel and highly relevant to clinical practice. It should be noted that we studied a healthy rat model. Kidney insults to those with baseline compromise may be even more significant.

Our findings represent the first preclinical analysis to our knowledge on the safety of vancomycin loading doses using specific biomarkers for kidney function and injury. In this study, rats received allometrically scaled doses of vancomycin corresponding to a guideline-recommended loading dose of 35 mg/kg/dose in humans, and a maintenance dose of 15 mg/kg/dose in humans. The finding of an immediate, significant decline in GFR and elevation of urinary KIM-1 after receipt of a vancomycin loading dose is highly relevant. Many patients who qualify for vancomycin loading doses are critically ill and have pre-existing renal dysfunction. Due to the known impact of AKI on inpatient mortality, all efforts should be made to characterize and subsequently minimize potential nephrotoxic dosing (20, 21). Our pre-clinical findings require further investigation and validation in other animal models and humans. However at minimum, these preliminary results provide evidence that vancomycin loading doses cause early kidney injury and loss of function in an animal model. Thus, loading doses should only be considered in those patients where expected gains are anticipated to outweigh the potential for increased kidney injury. The recently updated guidelines for therapeutic monitoring of vancomycin in serious MRSA infections recommend use of loading doses in critically ill patients with suspected or documented serious MRSA infections in order to improve efficacy (2). Clinical studies underpinning this recommendation are limited by their retrospective nature, use of previous trough goals, and use of the non-specific serum creatinine as a marker for both kidney function and injury (3, 4, 13-15). While multiple studies have found that administration of a loading dose results in higher serum vancomycin levels and faster achievement of designated PKPD targets for efficacy, few have assessed the potential nephrotoxicity associated with these doses (4, 14, 16). A recent retrospective cohort study by Flannery, et al. investigated the efficacy and safety of vancomycin loading doses in critically ill patients with MRSA infections (17). In terms of nephrotoxicity, this study found no difference in AKI rates among patients who received a vancomycin load and those who did not. However, serum creatinine was used to assess kidney injury, which is well-known to be an insensitive and delayed marker of AKI (18, 19).

In our investigation of variable vancomycin doses with an AKI-positive control, we found that receipt of intraperitoneal folic acid (AKI-positive control group) resulted in substantial kidney injury with near complete functional loss. This was characterized by a 45% increase in urinary output and 88% decrease in GFR compared to baseline, over the course of the study). Folic acid, a methodologic positive control known to induce oxidative damage in the proximal tubular epithelial cells, (22) was demonstrated to induce a much more significant kidney injury and loss of function when compared to rats that received low and high doses of vancomycin. These results are useful for understanding the relative damage that vancomycin can do. The urinary injury biomarker KIM-1 was most elevated in the AKI-positive control group on all experimental days, except for day 4 where KIM-1 was most elevated in the high dose vancomycin group. This finding shows that folic acid induced a greater degree of acute proximal tubular damage, compared to vancomycin at the administered doses. The finding of a KIM-1 elevation on day 4 among rats in the high dose vancomycin group is notable and may reflect cumulative kidney injury from repeated dosing. At the level of the individual rat, 3 of the 6 rats experienced an increase in urinary KIM-1 from day 3 to day 4. Measured urinary KIM-1 in the low and high dose vancomycin groups reached a maximum of 28 and 79%, respectively (on day 2 for all groups), of the KIM-1 levels seen in the AKI-positive control group. Clusterin was most elevated among rats in the high vancomycin dose group, second highest among rats in the AKI-positive control group, while rats in the low vancomycin dose group experienced only mild elevations from baseline. Clusterin in the low and high dose vancomycin groups was 39 and 108%, respectively, of the clusterin levels seen in the AKI-positive control. The observed trends in urinary clusterin may relate to its variety of proposed functions, including both cellular damage and repair processes (23, 24). When paired with the GFR findings for this experimental arm, we see that rats in the high vancomycin group experienced a mean GFR decrease of 43% from baseline to day 1, while rats in the AKI-positive control group experienced a mean GFR decrease of 89% over the same time period. Consequently, the observation of the highest degree of urinary clusterin expression among the high vancomycin dose group rats is not reflective of the greatest degree of kidney injury or loss of function. Rather, rats in the high vancomycin dose group may have experienced comparatively less kidney injury, followed by recovery, when compared to rats in the AKI-positive control group. Similar to the results from the loading dose arm, KIM-1 was the biomarker that most correlated with declining GFR over the course of the experiment. Our results help to further develop and characterize a translational rat model, specifically in reference to severe drug induced kidney injury and toxicodynamics of antibiotics in an animal model (6, 8, 11, 12, 25).

There are several considerations for our study. First, due to technical difficulty, two animals did not have adequate plasma collected on one day for each animal. These animals were still included in our analysis because the statistical methodology we utilized is flexible and does not require dropping the data (such as with analysis of variance [ANOVA] methods). Both animals provided adequate plasma data on all other experimental days, in addition to daily urine and terminal kidney samples. Second, we measured total urinary volumes in the rats, thereby capturing the total amounts of excreted biomarkers (versus quantification of spot samples) and standardized these to 24 hour excretion amounts. Although our findings do not change significantly when analyzed as 24-hour concentrations of biomarkers; the more exact methods available for laboratory studies should be considered when comparing to clinical studies where 24-hour urine collection is logistically more difficult. Future preclinical studies should evaluate GFR and urinary biomarkers over a longer period of time (i.e. 7 to 14 days) in order to understand if kidney function and injury can recover to baseline levels after receipt of a vancomycin loading dose. Clinical studies evaluating the safety of vancomycin loading doses should be conducted in a prospective manner, employing improved markers of kidney function and injury such as the ones utilized in this study.

## Materials and Methods

### Experimental design and animals

The experimental methods were similar to those we have previously reported (7-10). All experiments were conducted at Midwestern University in Downers Grove, Illinois, in compliance with the National Institutes of Health Guide for the Care and Use of Laboratory Animals (26) and were approved under the Institutional Animal Care and Use Committee protocol #3151. In brief, male Sprague-Dawley rats (n=34; age: 8 to 10 weeks; mean weight: 274.9 g) were housed in a light- and temperature-controlled room for the duration of the study and allowed free access to water and food (Supplemental figures 1 and 2). All animals were placed in metabolic cages (Lab Products, Inc., Aberdeen, MD, USA) for 24-hour urine collection, starting prior to dosing (day 0) and sampled every day for a period of 4 days. Animals were assigned to two experimental arms: 1) investigation of the impact of vancomycin loading doses and 2) investigation of variable vancomycin doses with an AKI-positive control. In the first arm, rats were assigned to one of two treatment groups in which they received either VAN 220 mg/kg intravenously over 2 minutes followed by VAN 100 mg/kg 12 hours later (n=9) [VAN loading dose group], or VAN 100 mg/kg intravenously over 2 minutes followed by VAN 100 mg/kg 12 hours later (n=7) [no VAN loading dose group] (Supplemental figure 3). No additional doses were given for the remainder of the study days (i.e. through day 4). In the second arm, animals were assigned to one of three treatment groups in which they received either intraperitoneal folic acid 250 mg/kg on day 1, followed by folic acid 100 mg/kg on days 2-4 (n=6), VAN 150 mg/kg/day intravenously over 2 minutes (n=6), or VAN 250 mg/kg/day intravenously over 2 minutes (n=6) (Supplemental figure 4). Rats in both VAN groups also received intraperitoneal injections of normal saline at equivalent volumes to the folic acid group. Following study drug dosing on each day, all animals received iohexol 51.8mg/day intravenously over 1 minute. Timed plasma samples were also drawn on each experimental day. Vancomycin doses were selected to approximate human doses allometrically scaled for the rat (100 mg/kg in rat ≈ 15 mg/kg in human, 220 mg/kg in rat ≈ 35 mg/kg in human). Following completion of the dosing protocol, all rats were euthanized and underwent nephrectomies.

### Chemicals and reagents

Rats were administered clinical grade vancomycin (Lot number: 167973; Fresenius Kabi, Lake Zurich, IL, USA), folic acid (Lot number: WXBD4723V; Sigma-Aldrich, St. Louis, MO, USA), iohexol [Omnipaque] (Lot number: 15025174; GE Healthcare Inc., Marlborough, MA, USA), and normal saline for injection (Hospira, Lake Forest, IL, USA). Vancomycin was prepared by weighing and dissolving the powder in normal saline to achieve a final concentration of 100 mg/mL. Folic acid was prepared by weighing and dissolving the powder in 0.3 mM sodium bicarbonate to achieve a final concentration of 50 mg/mL. Analytical grade iohexol (Lot number: LRAC5648; Sigma-Aldrich, St. Louis, MO, USA) and iohexol-d5 (Lot number: 28540; Cayman Chemical, Ann Arbor, MI, USA) were used for liquid-chromatography with tandem mass spectrometry analyses of plasma samples.

### Blood, urine, and kidney sampling

Double jugular vein catheters were surgically implanted 72 h prior to protocol initiation. One catheter was dedicated to blood sample draws, while drug dosing occurred via the other catheter. Blood samples were obtained from the catheter at pre-specified timepoints (0, 30, 60, and 240 minutes after iohexol dosing). Each sample (0.2 mL/aliquot) was replaced with an equivalent volume of normal saline (NS) for maintenance of euvolemia. Blood samples were prepared as plasma with EDTA (Sigma-Aldrich Chemical Company, Milwaukee WI, USA) and centrifuged at 3000 g for 10 minutes (Thermo Fisher Scientific, Waltham, MA, USA). Supernatants were collected and frozen at -80℃ until time of batch analysis with liquid chromatography tandem mass spectrometry (LCMS).

Urine samples were collected, and volume was measured starting from day 0. Urine collections at the day 1-PM and day 4 timepoints represented 12 and 4-hour urine collections, respectively. Measured volumes were multiplied in order to project full 24-hour urine production volume. All other timepoints (i.e. day 0, day 1-AM, day 2, and day 3) represented 24-hour urine collections. Samples were centrifuged at 400 g for 5 minutes at 4°C, and the resulting supernatant was collected, aliquoted, and stored at-80℃ until batch analysis of renal biomarkers. Following completion of the dosing protocol, rats were sacrificed.

### GFR measurement

GFR was assessed by iohexol clearance, as described in the model building section. Iohexol was administered intravenously as an undiluted solution on each dosing day, after administration of the study drug treatment. Rats received once-daily doses of iohexol 51.8mg/0.22 mL, given over 1 minute, on experimental days 0 through 4.

### Calibration curves in rat plasma

Stock solutions of iohexol and creatinine were prepared at a concentration of 1 mg/mL, while iohexol-d5, and creatinine-d3 were prepared at a concentration of 100 µg/mL. All drugs were dissolved in purified water. An iohexol standard curve was created by diluting stock solution with water to obtain concentrations between 0.5 and 100 µg/mL. Iohexol-d5 was used as the internal standard and was added to each sample to obtain a final concentration of 10 µg/mL. For the creatinine standard curve, stock solution was diluted with water, plasma was then spiked to obtain concentrations between 3.1 and 400 µg/mL. Creatinine-d3 was used as the internal standard and was added to each sample to obtain a final concentration of 10 µg/mL.

For each standard curve concentration, the iohexol or creatinine dilution (4 µL each) was added to 36 µL of blank rat plasma. Iohexol-d5 (4 µL), and creatinine-d3 (4 µL) were added to each standard curve concentration as internal standards. Each standard curve concentration was then mixed with 140 µL of 0.1% formic acid in methanol, vortexed, and centrifuged at 16,000 g for 10 minutes. Resulting supernatant was then transferred to LCMS vials for analysis.

### Sample preparation

For preparation of samples, 4 µL of iohexol-d5, and 4 µL of creatinine-d3 were added to 40 µL of sample rat plasma. Plasma samples at 30 and 60 minutes were diluted 1:10 with blank rat plasma. Each sample was then mixed with 136 µL of 0.1% formic acid in methanol, vortexed, and centrifuged at 16,000 g for 10 minutes. Resulting supernatant was then transferred to LCMS vials for analysis.

### LCMS methods

An Agilent 1260 series liquid chromatography system paired with 6420 triple quadrupole mass spectrometer (Agilent Technologies, Santa Clara, CA) was used to analyze plasma samples. Column temperature was maintained at 20°C. Mobile phase consisted of 0.1% formic acid in water (mobile phase A) and acetonitrile (mobile phase B) for both methods. For analysis of iohexol, a Poroshell 120 analytical column was used (2.7 µm, 100 × 3.0 mm, Part # 695975-302, Agilent Technologies). A gradient method was used to separate the analytes with the following solvent compositions: 5%[A]/95%[B] (0-4 mins), 97%[A]/3%[B] (4.1-7 mins) at a mobile phase flow rate of 0.6 mL/min. Multiple-reaction monitoring mode was used to analyte detection, and the transitions monitored were *m/z* 821.8 to 803.8 and *m/z* 826.8 to 808.8 for iohexol and iohexol-d5, respectively.

Creatinine was analyzed as previously reported (Pais et al., 2020). In brief, a Poroshell 120 analytical column was used (2.7 µm, 50 × 3.0 mm, Part # 699975-302, Agilent Technologies). A gradient method was used to separate the analytes and mobile phase flow rate was 0.6 mL/min. Multiple-reaction monitoring mode was used to analyte detection, and the transitions monitored were *m/z* 114.1 to 44.3 and *m/z* 117.1 to 89.2 for creatinine and creatinine-d3, respectively.

### Model building

In order to describe iohexol clearance, a two-compartment model was created for samples obtained in the post-distribution phase (Monolix 2021R1; Lixoft, Antony, France). Evaluated covariates included weight, log-transformed weight, and treatment group. To capture daily clearance changes, each experimental day was considered a separate occasion with clearance that could vary. Selection of the final model was based on the Akaike Information Criterion (AIC), between subject variability of the population estimates, goodness-of-fit plots for observed versus predicted, and the rule of parsimony. Empirical Bayes Estimates (i.e. individual Bayesian posteriors) were utilized for individual animal parameters.

### Determination of urinary biomarkers of AKI

Microsphere-based Luminex xMAP technology was used for the determination of urinary concentrations of KIM-1, clusterin, and OPN, as previously detailed (10, 12, 25). In brief, urine samples were allowed to thaw at ambient room temperature, aliquoted into 96-well plates, and mixed with Milliplex MAP rat kidney toxicity magnetic bead panel 1 (EMD Millipore Corporation, Charles, MO, USA). For each 96-well plate, a separate standard curve was prepared and run, per the manufacturer’s instructions. Results were analyzed and urinary biomarker concentrations were determined using five-parameter (linear and logarithmic scale) curve-fitting models (Milliplex Analyst 5.1; VigeneTech, Carlisle, MA, USA). Urinary biomarker concentrations were then normalized to 24-hour totals, based on individual measured urine volumes.

### Statistical analysis

A mixed-effects, restricted maximum likelihood estimation regression was used to compare urine output, mean weight loss, GFR, and urinary biomarkers among the treatment groups, with repeated measures occurring over days; measures were repeated at the level of the individual rat (Stata version 16.1, StataCorp LLC, College Station, TX, USA), thus correcting for multiple comparisons. Spearman’s rank correlation coefficient with a Bonferroni correction was used to assess correlations between kidney injury (e.g. KIM-1, clusterin, OPN) and function (e.g. GFR) by treatment day. Urinary biomarker amounts were log-transformed as needed for relationship exploration. All tests conducted were two-tailed, with an *a priori* level of statistical significance set at α = 0.05.

## Acknowledgements

The research reported in this publication was supported in part by the National Institute of Allergy and Infectious Diseases under award number R21-AI149026 (author MS), and a Pilot Research Award from Midwestern University (author JC). The content is solely the responsibility of the authors and does not necessarily represent the official views of the National Institutes of Health. MS reports ongoing research contracts with Nevakar and SuperTrans Medical as well as having filed patent US10688195B2. All other authors have no other related conflicts of interest to declare.

## Supplemental figures

(see supplemental document)

